# Systems-level immunomonitoring using self-sampled capillary blood

**DOI:** 10.1101/694521

**Authors:** Vijay Sai Josyula, Tadepally Lakshmikanth, Jaromir Mikes, Yang Chen, Petter Brodin

## Abstract

Comprehensive profiling of the human immune system in patients with cancer, autoimmune disease and during infections are providing valuable information that help us understand disease states and discriminate productive from inefficient immune responses and identify possible targets for immune modulation. Recent technical advances now allow for all immune cell populations and hundreds of plasma proteins to be detected using small volume blood samples. To democratize such systems-immunological analyses, further simplified blood sampling and preservation will be important. Here we describe that blood obtained via a nearly painless self-sampling device of 100 microliter of capillary blood that is preserved and frozen, can simplify systems-level immunomonitoring studies.

---

Systems immunology involves simultaneous analyses of all immune system components and their inter-relationship in health and disease. Such analyses are revealing important patterns, previously not visible using more reductionist approaches involving individual cell populations and proteins^1^. For example, we have learned in recent years that human immune systems are predominantly shaped by by non-heritable factors such as Cytomegalovirus^2–5^, and that immune systems diverge with age as environmental exposures accumulate. We are also beginning to learn that baseline immune system states can be predictive of vaccine responses^3,6^, and response to immunomodulatory treatments^7^. During an immune response, for example in the context of immunotherapy of cancer, signatures of immune cell changes can predict clinical outcome^8^. Finally, by longitudinal monitoring of healthy individuals we are learning how newborn immune systems are shaped early in life^9^, and how immunological changes associated with aging manifest itself and affect disease risks in the elderly^10^.

We have recently described that using stabilized and frozen whole blood, rather than the more commonly used viable peripheral blood mononuclear cells for immunomonitoring, offers advantages thanks to lower technical variation^11^. and Using preserved whole blood also allow for much smaller blood volumes to be used^9^. In parallel with this, Blicharz and colleagues have developed a microneedle-device for self-sampling of capillary blood that is meant to simplify blood sampling and mitigate the fear of needles and simplify clinical blood testing^12^. The system is virtually pain-free due to the use of microneedles that only sample shallow capillaries in the skin (Figure 1a-b). Here we show, that patient self-sampling of capillary blood using TAP (7sense Bio, Medford, MA, USA), in combination with whole blood stabilizer (Whole blood processing kit, Cytodelics AB, Stockholm, Sweden), and an optimized cell processing protocols^13^, allows for complete systems-level immunomonitoring. This will enable samples to be collected at home by patients without having to visit a clinic, allowing more frequent sampling and more careful monitoring of their immune systems. We show, using Mass cytometry that system-level immune profiles are comparable to those obtained via traditional venipuncture sampling protocols, and that the blood volumes obtained by self-sampling devices are enough to capture comprehensive immune system states across all relevant immune cell populations.

**Figure 1.**
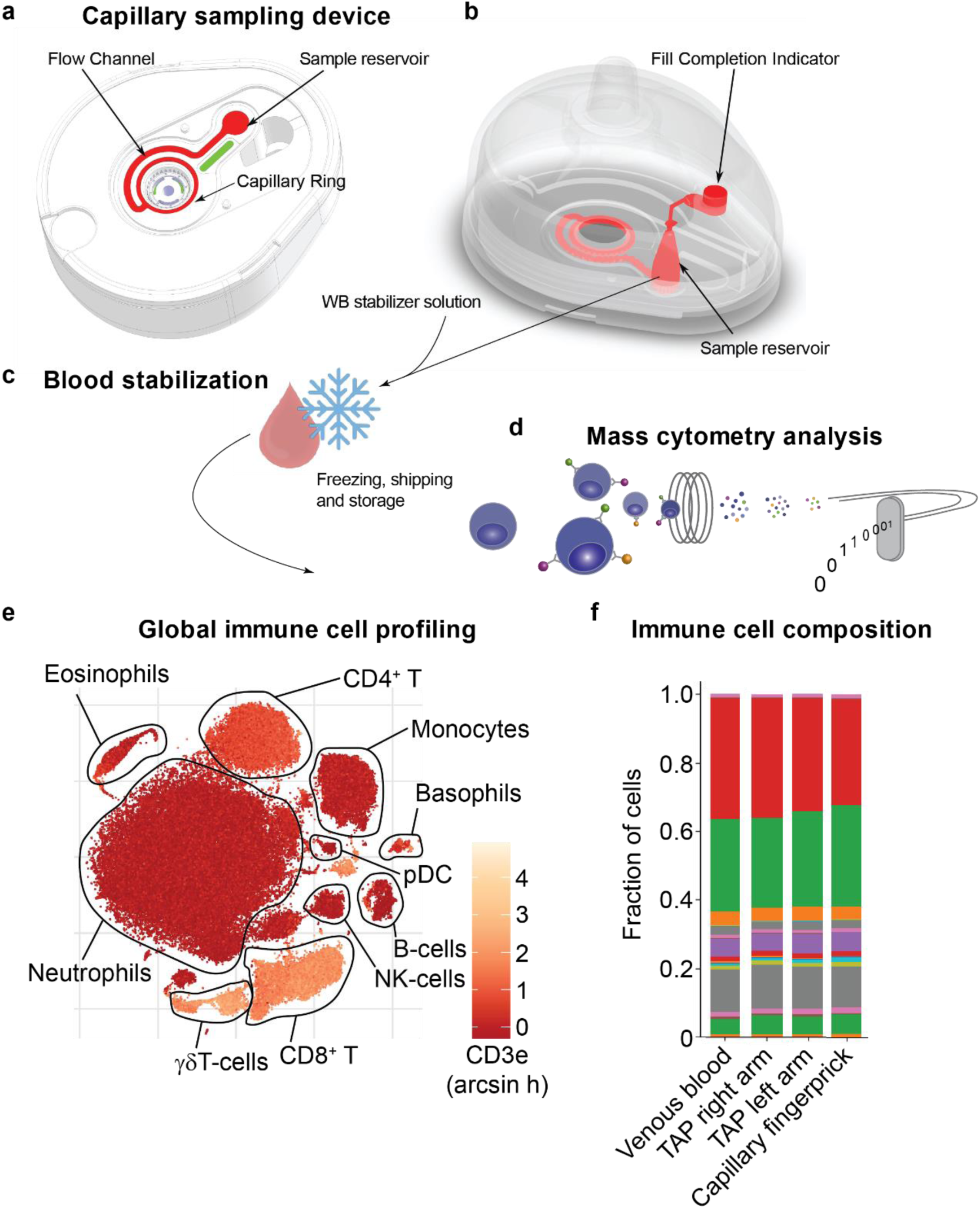
Systems-level immunology via a self-sampling device. a-b) Device for self-use capillary sampling, c) Stabilization of whole blood and freezing prior to Mass cytometry analysis. e) Example tSNE of a blood sample collected via TAP-capillary device, and f) relative proportions of major immune cell populations in TAP-sampled, finger prick and venipuncture sampled blood.

To test the self-sampling system for immunomonitoring we allowed non-experts to sample themselves using the TAP-device in both of their upper arm and directly after this, they were also subject to traditional venipuncture as well as a capillary finger-prick sampling using traditional lancet. All the samples from these localities were preserved using a whole-blood stabilizer solution and frozen at −80 C and transported to the laboratory for analysis (Figure 1c). From our previous testing we know that also −20 C freezing of whole blood is possible for a limited amount of time, allowing subjects to freeze samples at home. The blood samples were barcoded together^14^, stained with a 48-parameter panel targeting markers in all major immune cell populations. The samples were subsequently acquired by Mass cytometry and canonical cell populations identified using a machine learning approach (Chen et al, manuscript in preparation). The results show that cell frequencies differ among the two donors as expected from the known variation among healthy human immune systems^15^ (Figure 1c). We conclude from this result that self-sampling using the nearly painless TAP device, in combination with whole blood stabilization and freezing can enable more broad application of systems-level immunomonitoring by allowing patients to sample themselves at home. Other important implications of this work are that patients soon will be able to sample themselves frequently in the comfort of their own home, without risking significant blood loss over time thanks to the small volumes collected with each sample, possibly increasing the temporal resolution in many immunomonitoring studies.

All in all, we have shown here that the combination of two recently developed methods for improved blood sampling now allow for advanced, systems-level immunomonitoring using minimal samples of capillary blood collected by patients themselves, with broad implications for studies in a range of patients with immune mediated disease, infections and cancer, treated with immunotherapy to name a few.

## Methods

### Blood sample collection and processing

Blood samples (∼100μl) were obtained from two healthy donors from the following sites using the following devices – a) Capillary blood collection (TAP device, 7sense bio) – blood was collected from both upper left and right arms; b) Venipuncture (vacutainer blood collection tube and needle) – venous blood was drawn from the middle cubital vein in the arm; c) Fingerstick (Lancet and microvette) – blood was drawn from one of the fingertips. Each blood sample was mixed 1:1 with a blood stabilizer solution (Cytodelics AB, Stockholm), incubated for 10 min at room temperature followed by freezing at −80 C. At the time of experimentation, blood samples were thawed and fixed/lysed using Fix/Lyse buffer (Cytodelics AB) as per the manufacturer’s recommendations.

### Sample multiplexing and staining

Blood samples post fix/lysis were resuspended in Wash # 2 buffer (Cytodelics) and barcoded using 20-Plex Pd Barcoding kit (Fluidigm). Samples were barcoded using the principles reported elsewhere^14^. A Bravo liquid handling automation platform^13^ (Agilent technologies) was used to barcode and pool samples. Briefly, each barcode was resuspended in 100ul of 1x barcode perm buffer. The cells that were resuspended in Wash # buffer and centrifuged at 500g for 5 min followed by 2 washes with 1x barcode perm buffer. Following washes, samples were resuspended in 100ul of 1x barcode perm buffer and to this the reconstituted barcodes were added, incubated for 30 min at room temperature followed by 2 washes with CyFACS buffer (PBS 1x with 0.1% BSA, 2mM EDTA and 0.05% Na-Azide) and pooled according to the sample schema.

The pooled sample batches were washed and FcR blocked, following which they were stained with a cocktail of metal-conjugated antibodies targeting surface markers. Cells were incubated at 4 C for 30 min, washed twice using CyFACS buffer at 500g for 5 min and fixed overnight in 4% PFA (diluted in PBS) before mass cytometry analysis.

### Antibodies for Mass cytometry

The monoclonal antibodies used in this study are listed in Table 1. Pre-conjugated antibodies were bought from Fluidigm if the metals were unavailable in their pure form. Other metals that were available in a pure form were purchased and conjugated to purified antibodies (obtained in carrier/protein-free buffer) using MAXPAR X8 polymer conjugation kit (Fluidigm) by following the manufacturer’s protocol. The antibody concentrations were measured using NanoDrop 2000 spectrometer (Thermo Fischer Scientific) at 280nm before and after conjugation. The conjugated antibodies were then diluted 1:1 using Protein Stabilizer PBS (Candor Bioscience GmbH).

**Table 1.** Antibodies used for mass cytometry analysis.

| **Metal tag** | **Marker** | **Clone** | **Company\*** |
| --- | --- | --- | --- |
| 89Y | CD45 | HI30 | Fluidigm |
| 102Pd | Barcode | Cell-ID™ 20-Plex Pd Barcoding Kit | Fluidigm |
| 104Pd | Barcode |  |  |
| 105Pd | Barcode |  |  |
| 106Pd | Barcode |  |  |
| 108Pd | Barcode |  |  |
| 110Pd | Barcode |  |  |
| 113In | CD57 | HCD57 | BioLegend |
| 115In | HLA-A, B, C | W6/32 | BioLegend |
| 141Pr | CD49d | 9F10 | Fluidigm |
| 142Nd | CD19 | HIB19 | Fluidigm |
| 143Nd | CD5 | UCHT2 | BioLegend |
| 144Nd | CD16 | 3G8 | BioLegend |
| 145Nd | CD4 | RPA-T4 | BioLegend |
| 146Nd | CD8a | SK1 | BioLegend |
| 147Sm | CD11c | Bu15 | Fluidigm |
| 148Nd | CD31 | WM59 | BioLegend |
| 149Sm | CD25 | 2A3 | Fluidigm |
| 150Nd | CD64 | 10.1 | BioLegend |
| 151Eu | CD123 | 6H6 | BioLegend |
| 152Sm | TCR$\gamma\delta$ | 5A6.E9 | Fischer Scientific |
| 153Eu | Siglec-8 | 837535 | R&D Systems |
| 154Sm | CD3e | UCHT1 | BioLegend |
| 155Gd | CD33 | WM53 | BioLegend |
| 156Gd | CD26 | BA5b | BioLegend |
| 157Gd | CD9 | SN4 C3-3A2 | eBiosciences |
| 158Gd | CD34 | 581 | BioLegend |
| 159Tb | CD22 | HIB22 | BioLegend |
| 160Gd | CD14 | M5E2 | BioLegend |
| 161Dy | CD161 | HP-3G10 | BioLegend |
| 162Dy | CD29 | TS2/16 | BioLegend |
| 163Dy | HLA-DR | L243 | BioLegend |
| 164Dy | CD44 | BJ18 | BioLegend |
| 165Ho | CD127 | A019D5 | Fluidigm |
| 166Er | CD24 | ML5 | BioLegend |
| 167Er | CD27 | L128 | Fluidigm |
| 168Er | CD38 | HIT2 | BioLegend |
| 169Tm | CD45RA | HI100 | Fluidigm |
| 170Er | CD20 | 2H7 | BioLegend |
| 171Yb | CD7 | CD7-6B7 | BioLegend |
| 172Yb | IgD | IA6-2 | BioLegend |
| 173Yb | CD56 | NCAM16.2 | BD |
| 174Yb | CD99 | HCD99 | BioLegend |
| 175Lu | CD15 | W6D3 | BioLegend |
| 176Yb | CD39 | A1 | BioLegend |
| 191Ir | DNA-Ir | Cell-ID™ Intercalator-Ir | Fluidigm |
| 193Ir |  |  |  |
| 209Bi | CD11b | Mac-1 | Fluidigm |

*\* All antibodies other than those purchased from Fluidigm have been coupled in-house*

### Sample acquisition by CyTOF

To cells fixed in PFA, Cell-ID Intercalator-Ir (191Ir/193Ir) (Fluidigm) diluted 1:1000 (stock 125 μM) was added and incubated for 20 min at room temperature. Cells were washed twice with CyFACS buffer, followed by PBS, and milli Q water and resuspended in milli Q water. Cells were counted and filtered through a 35µm nylon mesh, diluted to 750,000 cells/ml by mixing 0.1X times with EQ™ four element calibration beads (Fluidigm) in Milli-Q water and acquired at a rate of 300-500 cells/s using a CyTOF2 (Fluidigm) mass cytometer, CyTOF software version 6.0.626 with noise reduction, a lower convolution threshold of 200, event length limits of 10-150 pushes, a sigma value of 3, and flow rate of 0.045 ml/min. All raw FCS-files are available https://flowrepository.org/, ID: FR-FCM-Z25W.

### Data analysis

#### Preprocessing of data

The samples were debarcoded and relative proportion of the major cell types in each sample was obtained using an automated classification algorithm trained on manually gated training datasets (Chen et al, manuscript). The output file is a single-cell table with cells in rows and markers in columns and an annotation column showing population label according to the learning algorithm.

#### Statistics

The relative proportions of the cell populations of each sample was plotted as a stacked bar plot using matplotlib and the pandas libraries in python version 3.0.

#### Data visualization

Single-cell data were embedded by tSNE, using the R-implementation, rtsne after scaling all markers to unit variance. This was done using R version 3.6.0 (https://www.r-project.org/). An Aitchinson’s distance matrix was calculated between all samples using their relative cell proportions as input using the aDist function. Multidimensional scaling coordinates were obtained using the cmdscale function and the resultant 2D data visualized as a scatter plot using the ggplot2 package.

## Acknowledgements

V.S.J performed experiments with the help from T.LK. The study was planned by P.B, T.LK, J.M and Y.C provided data analysis support.

